# Vertical distribution of *Phytophthora agathidicida* oospore DNA in kauri forest soils: Implications for optimised sampling and disease monitoring

**DOI:** 10.64898/2026.03.26.714588

**Authors:** Jade T. T. Palmer, Emily M.G. Hocking, Monica L. Gerth

## Abstract

*Phytophthora* species are globally significant soilborne oomycetes responsible for widespread ecosystem decline. Standard soil sampling protocols, originally developed for qualitative baiting assays, typically require collecting substantial soil volumes in order to capture viable propagules. While effective for culture-based detection, these protocols are labour-intensive and can damage the shallow root systems of sensitive host species such as New Zealand kauri (*Agathis australis*).

*Phytophthora agathidicida* (PA), the pathogen associated with kauri dieback disease, is routinely surveyed using these methods. However, quantitative data describing the vertical distribution of PA in natural forest soils are lacking. Consequently, it remains unclear whether extensive depth sampling is necessary to ensure consistent molecular detection.

In this study, we applied a quantitative oospore DNA (oDNA) qPCR assay to characterise the fine-scale vertical distribution of PA across four soil depth increments (0–5, 5–10, 10–15, 15–20 cm) from 12 kauri trees representing a range of disease stages. Results revealed distinct vertical stratification, with PA DNA concentrations peaking within the upper 0–10 cm of soil in non-symptomatic and possibly symptomatic trees. In symptomatic trees, the absolute peak occasionally reached 10–15 cm, while pathogen signals remained consistently detectable within the top 10 cm. Field validation from an additional eight trees confirmed that targeted 0–10 cm “shallow” sampling yielded higher PA concentrations than deeper sampling protocols.

These findings provide a data-driven basis for refining soil sampling strategies, enabling more sensitive molecular detection while minimising disturbance and logistical effort in fragile ecosystems.

**IMPORTANCE:** *Phytophthora* species are among the most destructive soilborne pathogens globally, requiring robust diagnostic protocols for both agricultural and conservation settings. Traditional sampling frameworks were established to meet the biological requirements of baiting assays, which often necessitate collecting large soil volumes from broad depth profiles to ensure the capture of viable, infectious propagules. However, these extensive requirements are labour-intensive and can cause significant soil disturbance in sensitive forest ecosystems.

Using *P. agathidicida* as a model, this study provides a high-resolution quantitative assessment of how pathogen DNA is distributed vertically across different disease stages. We demonstrate that while absolute peak abundance can shift within the 0–15 cm range as infection progresses, the pathogen signal remains consistently detectable within the top 10 cm. This evidence-based approach suggests that targeted, shallow sampling enhances sensitivity by reducing signal dilution, offering a lower-impact path for monitoring soilborne oomycetes worldwide.

## INTRODUCTION

*Phytophthora* species are among the most destructive soilborne oomycetes worldwide, causing severe agricultural losses and widespread ecosystem decline (1-4). Traditional sampling protocols for these pathogens have been standardised around the requirements of baiting methods. Because these qualitative assays depend on the successful—and often inconsistent—germination of propagules to produce motile zoospores, they typically require large soil volumes collected from broad depth profiles to ensure pathogen capture. However, molecular diagnostics that utilise smaller soil volumes and enable quantification of pathogen load create a critical opportunity to refine legacy sampling volumes and depths without compromising detection sensitivity.

A critical example is *Phytophthora agathidicida* (PA), which causes widespread dieback of kauri (*Agathis australis*) trees in New Zealand’s northern forests (5-7). Kauri possess extensive, shallow, and highly sensitive fine-root systems concentrated in surface soils, and soil disturbance near these roots is strongly discouraged (8). Sampling protocols for PA have recommended collecting composite soil samples comprising subsamples taken from several points (typically 4–8) around a tree, to a depth of approximately 15–20 cm, yielding a total soil volume of roughly 1 kg per sample (9, 10). These methods are labour-intensive, disturb fine kauri roots, and require considerable effort for sample storage, transport, and disposal.

There is limited empirical evidence underpinning current PA soil sampling protocols. One study examined soil from two symptomatic kauri trees using baiting to assess depth distribution (11). Detection was successful to depths of 14 cm beneath one tree and 20 cm beneath the second (11). However, the small sample size and the qualitative nature of baiting mean little is known about the abundance of *P. agathidicida* beyond its potential presence in the upper 0–20 cm. Studies of other *Phytophthora* species generally report the highest recovery from upper soil layers (0–10 cm). However, results vary among species and depend on factors such as host root distribution and environmental context (*e*.*g*., agricultural fields, forests, nurseries) (12-15). Sampling depths vary among studies, with some extending to 30 cm (16).

The development of a quantitative qPCR assay for *P. agathidicida* oospore DNA (oDNA) enables the direct measurement of pathogen abundance from as little as 10–15 g of soil (17, 18). This shift toward smaller sample volumes necessitates a re-evaluation of optimal sampling depth. By using qPCR to map the vertical distribution of PA, we can optimise sampling protocols to maximise detection sensitivity while minimising environmental impact and logistical burden.

Here, we present the first quantitative assessment of *P. agathidicida* oospore DNA distribution across soil depths in kauri forests, examining 20 trees across two sites and multiple disease stages. These data inform evidence-based recommendations for revised soil sampling protocols and demonstrate how quantitative molecular approaches can improve *Phytophthora* diagnostics and monitoring in sensitive forest ecosystems.

## MATERIALS AND METHODS

### Soil Core Sampling: Detailed Depth Profile Analysis

Soil samples were collected from twelve kauri trees at Kauri Mountain, Whangārei Heads, Northland, New Zealand. Trees encompassed a range of health statuses (*i*.*e*. non-symptomatic, potentially symptomatic and symptomatic) to capture potential variation in pathogen load. Symptom status was scored in the field by an experienced observer using standard visual assessment criteria (*i*.*e*. canopy condition and presence of lesions).

At each tree, four soil cores were collected at approximately 1 m distances from the trunk in the uphill, downhill, left, and right directions. Each core was subdivided into four depth fractions: 0–5 cm, 5–10 cm, 10–15 cm, and 15–20 cm. Sampling points were adjusted as necessary to avoid roots, rocks, or other obstructions, and alternative cores were taken within 30 cm of the original sampling location if the target depth could not be reached. When the upper 10 cm was too loose or fibrous for core sampling, the 0–5 cm and 5–10 cm layers were collected with a trowel, and the corer was used for the lower sections. All sampling equipment was disinfected with bleach wipes between each core and twice between trees to minimise the risk of cross-contamination.

Soil samples were stored at 4 °C until extraction, and all samples were processed within seven days of collection. Soil collected at each depth for each tree was pooled and hand-homogenised, yielding four composite samples per tree and 48 samples in total. All sample processing was conducted in a sterile laminar-flow hood, and laboratory surfaces and equipment were regularly decontaminated with bleach and a DNase-based reagent throughout the extraction procedure.

### Trowel-Based Soil Sampling: Comparing Depths in Field Conditions

In addition to core sampling, trowel-based sampling was conducted under field-realistic conditions to compare two depths: approximately 10 cm deep (“shallow”) or up to 20 cm (“deep”).

The Te Roroa Kauri Ora team collected eight soil samples around each of eight kauri trees; we requested that they sample trees with visible symptoms for this comparison. Samples at each depth were combined and homogenised in the lab into one composite sample per tree. This produced 16 composite samples total (eight shallow, eight deep) for oDNA-qPCR analysis.

### DNA Extraction and Storage

All 64 soil samples were processed using the oDNA method as previously described (17, 18). The only modification to the original protocol was in the final elution and storage steps. DNA was eluted in a low-EDTA TE buffer (10 mM Tris-HCl, 0.1 mM EDTA) rather than the manufacturer’s supplied buffer to ensure greater DNA stability during storage. Extracted DNA was stored at -20 °C in DNA LoBind tubes (Eppendorf), which help to reduce DNA adhesion to tube walls, thereby maximising DNA recovery for quantitative PCR (qPCR) and preserving sample integrity for potential downstream analyses.

### Quantitative PCR (qPCR)

All qPCR assays were performed using a QuantStudio3 Real-Time PCR System (Applied Biosystems). Each assay employed fluorescein amidite (FAM; Macrogen) as the fluorescent dye and Black Hole Quencher 1 (BHQ1; Macrogen) as the quencher.

The PA qPCR conditions were as follows: an initial denaturation at 95 °C for 10 minutes, followed by 40 cycles of 95 °C for 15 seconds and 58 °C for 60 seconds. Each qPCR reaction was set up in a 30 μL final volume containing 1× HOT FIREPol Probe Universal qPCR Mix, 400 nM of each primer, 250 nM of the TaqMan probe, and 3 μL of input DNA. The primers and probe used for quantification of PA were: PA-LTR-for 5′-ACGCGCTCTGTTTCTTTAGC-3′; PA-LTR-rev 5′-GCGGGCTTCCATTCAATTCA-3′; and PA-LTR-TaqMan probe, 5′-FAM-TAGTATGCGCTTTTGAGGAAGCGTAA-BHQ1-3′ (17, 18).

The genus-level *Phytophthora* qPCR conditions were as follows: an initial denaturation at 95 °C for 10 minutes, followed by 40 cycles of 95 °C for 15 seconds and 60 °C for 60 seconds. Each qPCR reaction was set up in a 30 μL final volume containing 1× HOT FIREPol Probe Universal qPCR Mix, 400 nM of each primer, 250 nM of the TaqMan probe, and 3 μL of input DNA. The primers and probe used for genus-level quantification of *Phytophthora* spp. were: PhyG-F2 5′-CGTGGGAATCATAATCCT-3′ and PhyG-Rb 5′-CAGATTATGAGCCTGATAAG-3′, along with the probe TrnM_PhyG_probe2 5′-ATRTTGTAGGTTCAARTCCTAYCATCAT-3′ (19).

For standard curve generation, genomic DNA was extracted from *P. agathidicida* isolate 3770. Mycelial mats were grown in the dark at 22°C in potato-dextrose broth (Difco) until they covered approximately 50% of the 90 mm Petri dish surface. The mats were harvested, washed with sterile water, dried using Kimwipes, and ground to a fine powder in liquid nitrogen using a mortar and pestle. DNA was extracted using the DNeasy Plant Mini Kit (Qiagen) as per the manufacturer’s protocol. DNA concentrations were quantified using the Qubit dsDNA High Sensitivity Assay kit.

Ten genomic DNA standards in a tenfold dilution series from 0.03 fg to 30,000,000 fg (30 ng) were prepared in DNA LoBind tubes (Eppendorf), with nine replicates per concentration. Cycle threshold (CT) values were plotted against log10-transformed DNA concentrations to construct the standard curve and calculate efficiency and R^2^ (Supplementary Figure 1A). The limits of detection and quantification were determined using previously developed algorithms (20)(Supplementary Figure 1B). Seven genomic DNA standards in a twofold dilution series from 0.16 fg to 60 fg were prepared in DNA LoBind tubes, with nine replicates per concentration. The coefficient of variation for the limit of quantification was set to 0.35, and both algorithms were set to ‘Best’, as recommended (20). The calculated limit of detection (LoD) was 0.7 fg, and the limit of quantification (LoQ) was 1 fg.

Subsequent qPCR experiments used seven standards from the most linear range of the standard curve, from 3 fg to 3,000,000 fg (3 ng), to determine the concentration of PA DNA in soil samples. All reactions were performed in triplicate, and final concentrations are expressed as fg of PA DNA per gram of soil (fg/g), accounting for the total elution volume of the DNA extract and the initial soil mass. Negative controls (distilled water) and a spiked inhibition control (3 fg spike of PA genomic DNA) were included in each run to monitor for contamination and potential PCR inhibition.

## RESULTS

### Relationships between PA abundance, depth, and symptom status

Twelve kauri trees of varying symptom status were sampled at four depth intervals (to 20 cm), yielding 48 total samples (**Figure 1**). Of these 12 sites, eight tested positive for PA DNA using the oDNA-qPCR method. Aggregated across these eight positive sites, mean PA DNA peaked at 200 fg/g soil (5–10 cm), declining to 25 fg/g in the deepest layer sampled (15–20 cm) (**Table 1)**.

**Figure 1.**
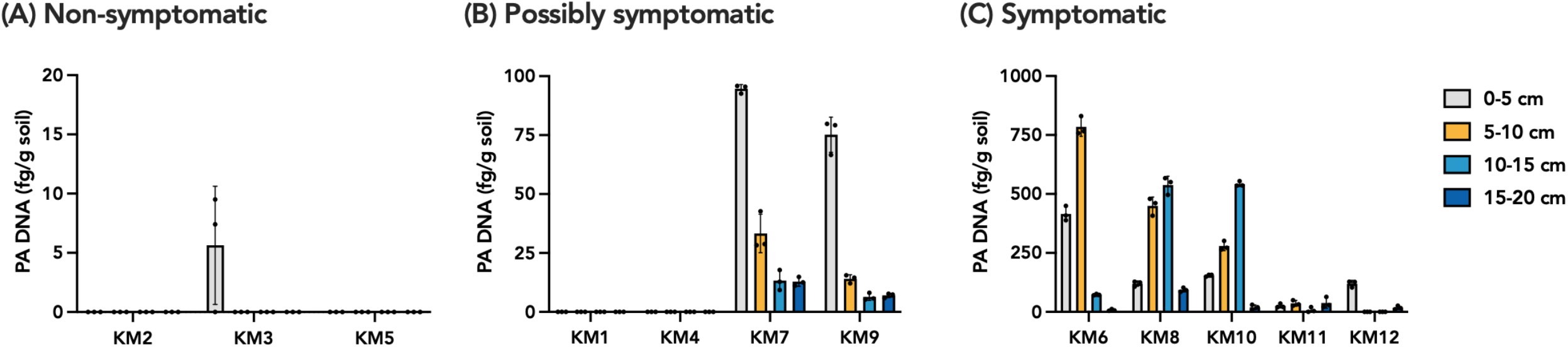
Distribution of *P. agathidicida* (PA) DNA at different depths beneath individual kauri trees with differing symptom status. Because PA DNA loads differed markedly across symptom categories, the y-axis scales differ between panels (A), (B), and (C). Soil cores were collected beneath kauri trees classified as (A) non-symptomatic, (B) possibly symptomatic, or (C) symptomatic, and sectioned into four depth intervals (0–5, 5–10, 10–15, and 15–20 cm; colour-coded as indicated in the key). Concentrations of PA DNA were quantified by qPCR and are expressed as fg PA DNA per g soil (mean ± standard deviation (n=3) for each tree (sample IDs KM1 to KM12).

**Table 1.** Vertical distribution of *Phytophthora agathidicida* (PA) DNA across all PA-positive kauri (n=8). Values represent the mean concentration per depth interval ± standard error.

| Depth | Average PA DNA (fg/g soil) |
| --- | --- |
| 0- 5 cm | 126 ± 6 |
| 5-10 cm | 200 ± 21 |
| 10-15 cm | 148 ± 31 |
| 15-20 cm | 25 ± 4 |

While the mean distribution showed a peak in the 5–10 cm layer, individual vertical profiles varied according to the observed disease status of the trees, revealing patterns relative to symptom severity:

- **Non-symptomatic trees** (n = 3): PA DNA was undetectable in two trees (KM2, KM5). The third (KM3) showed only low levels (∼5 fg/g soil) restricted to the surface 0–5 cm layer.
- **Possibly symptomatic trees** (n = 4): PA DNA was detected in two of the four trees, predominantly in the 0–5 cm layer at moderate levels (50-100 fg/g soil), with concentrations declining rapidly with depth.
- **Symptomatic trees** (n=5): PA was detected in all five symptomatic trees. These trees exhibited the broadest vertical spread of PA DNA. Elevated levels (∼100 fg/g soil or more) were detected across the top three layers (0–5, 5–10, and 10–15 cm), with markedly lower amounts at 15–20 cm.

Across symptom categories, PA DNA abundance increased progressively with disease severity. Among the symptomatic kauri, KM6, KM8, and KM10 showed the strongest signals (250–800 fg/g), while KM11 and KM12 had lower but clearly positive concentrations.

Analysis of these profiles showed that 7 of 8 positive trees reached their maximum DNA concentration within the top 10 cm of soil. Even in instances where maximum abundance occurred in deeper layers (*e*.*g*., KM10 at 10–15 cm), PA DNA in the upper horizons remained well above the detection threshold of 1 fg.

### Vertical distribution of *Phytophthora* spp. DNA

For comparison, the vertical distribution of *Phytophthora* spp. DNA was quantified using genus-level primers across the same depth profiles. While total concentrations varied significantly among individual trees—ranging from undetectable levels to >15,000 fg/g—the depth-related patterns were highly consistent across the study site. Regardless of a tree’s specific symptom status or total pathogen load, the highest abundance of *Phytophthora* spp. DNA in either of the two upper soil layers (0–5 cm and 5–10 cm) in 11 of the 12 trees sampled **(Figure 2)**. The sole exception was KM1, which had comparatively low levels (2–6 fg/g) in the bottom two layers (10–20 cm). Given that these levels represented a negligible fraction of the total site-wide pathogen load, they did not alter the overall trend. Collectively, these profiles reinforce that shallow sampling (0–10 cm) is optimal for detecting the vast majority of *Phytophthora* DNA in kauri forest soils.

**Figure 2.**
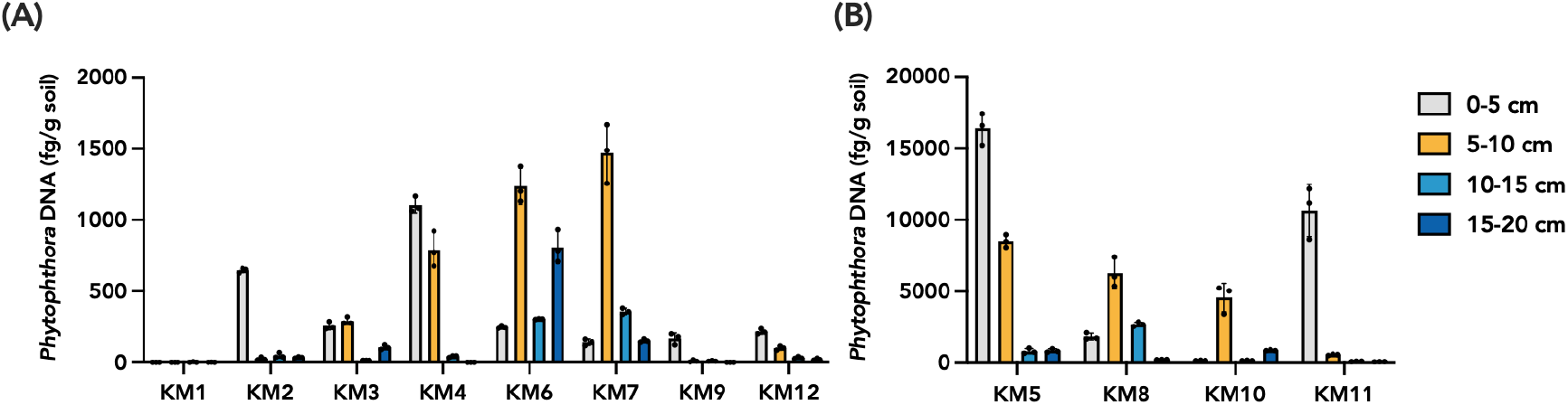
Distribution of *Phytophthora* spp. DNA at different soil depths. Because *Phytophthora* spp. DNA loads differed markedly across the sampled trees, the y-axis scales differ between panels (A) and (B). Soil cores were collected beneath 12 kauri trees classified and sectioned into four depth intervals (0–5, 5–10, 10–15, and 15–20 cm; colour-coded as indicated in the key). Concentrations of *Phytophthora* DNA were quantified by qPCR and are expressed as fg of DNA per g soil (mean ± standard deviation of three technical replicates) for each tree (sample IDs KM1 to KM12).

### Comparison of standard and modified shallow sampling protocols

The detailed depth profiles from the soil cores suggested that shallower sampling could effectively capture PA when present. To evaluate whether this finding could translate into routine field sampling, the Te Roroa Kauri Ora team collected soil samples around eight kauri trees considered potentially symptomatic. Two sampling methods were compared: the standard protocol, which combines soil from eight holes dug to approximately 20 cm depth into a single composite (“deep”) sample, and a modified “shallow” sampling approach, which collected soil from eight nearby holes dug only to about 10 cm depth.

Of the eight trees sampled, five tested positive for PA using both approaches; PA was not detected in the samples from TR1, TR4 and TR6. In all positive cases, the “shallow” soil samples contained higher concentrations of PA DNA than the corresponding “deep” samples (**Figure 3**).

**Figure 3.**
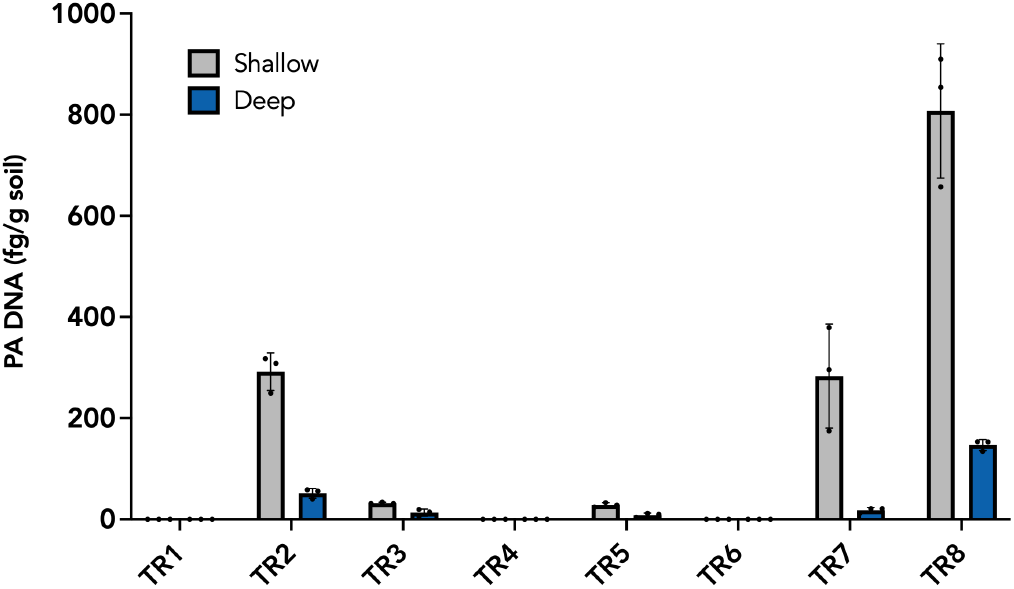
Quantification of *P. agathidicida* (PA) DNA at two soil sampling depths beneath kauri trees. Soil samples were collected beneath trees classified as possibly symptomatic at two depths: approximately 0–10 cm and 0–20 cm (colour-coded as indicated in the key). PA DNA concentrations were quantified by qPCR and are expressed as femtograms (fg) of PA DNA per gram of soil (mean ± range of three technical replicates) for each tree (sample IDs TR1–TR8). TR1, TR4, and TR6 were negative for PA DNA across both depths.

## DISCUSSION

To our knowledge, this is the first study to quantify PA oospore DNA at fine-scale depths around individual kauri trees. Our results reveal a distinct vertical stratification: the highest PA concentrations occurred in the upper 0–5 cm beneath non-symptomatic and possibly symptomatic trees, then shifted to the 5–15 cm layer beneath later-stage symptomatic trees. Because these patterns were consistently detectable within the top 10 cm across all positive cases, our data indicate that shallower sampling maximises detection sensitivity. In contrast, standard protocols that pool soil from 0–20 cm risk diluting pathogen signals, thereby reducing diagnostic performance.

Field testing by the Te Roroa Kauri Ora team further validated these findings. In a trial across eight trees, smaller, shallower cores (0–10 cm) yielded higher PA concentrations than standard deeper protocols, confirming improved sensitivity with reduced soil volumes. This consistency suggests that revising standard operating procedures to prioritise sampling at 0– 10 cm could enhance detection efficiency. Furthermore, while this study utilised molecular oDNA assays, these abundance profiles suggest that shallower sampling would likely benefit traditional soil baiting methods as well by concentrating the target inoculum.

While representing a single temporal snapshot, the data from trees at varying disease stages, coupled with the quantitative oDNA assay, also provide insights into disease dynamics. Our data support a model (**Figure 4**) in which, in early stages, PA is present at low levels, likely introduced at low concentrations through contaminated soil on footwear, equipment, or animals, or via water movement, with soils beneath apparently healthy kauri containing low levels of PA (<5 fg/g soil) at shallow depths. As infection progresses, pathogen replication within host tissues increases PA DNA concentrations, with early-stage or possibly symptomatic trees yielding up to ∼100 fg/g soil. In later stages, root dieback releases oospores into the surrounding soil; severely affected or dying trees generally contain more than 100 fg/g soil.

**Figure 4.**
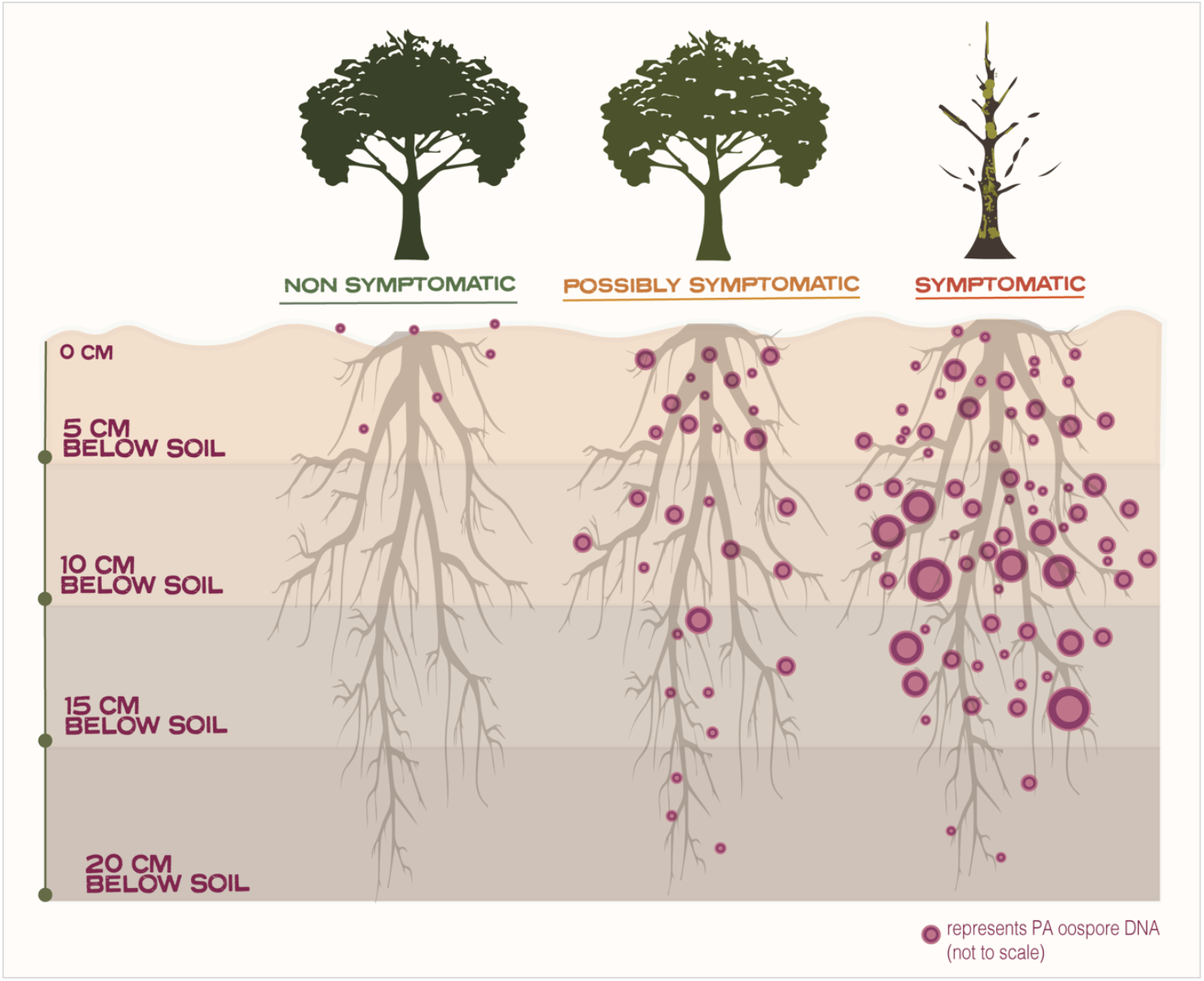
Schematic model of *P. agathidicida* (PA) oospore DNA vertical distribution patterns across kauri disease stages. This conceptual diagram illustrates general trends: PA DNA is typically shallowest in non-symptomatic trees, more widely distributed in possibly symptomatic trees, and peaks deeper in symptomatic trees while remaining detectable in upper soil layers. Note: Relative scales and patterns are illustrative.

These benchmarks may help interpret molecular detection results relative to disease severity. This raises the possibility of inferring infection stages from vertical profiles—a level of resolution previously unattainable with qualitative tools. By applying an oospore-targeted assay, this study demonstrates that molecular diagnostics can resolve fine-scale distribution patterns that mirror host-pathogen interactions. These benchmarks provide a quantitative framework for interpreting molecular detection results relative to field disease severity.

The relationships observed between inoculum load and disease stage underscore the specificity and robustness of the oDNA-qPCR assay. Overall, this approach offers distinct advantages over both traditional baiting and broad environmental DNA (eDNA) or metagenomic methods. Unlike baiting, which requires propagule germination—a biological process frequently confounded by dormancy or microbial competition—oDNA-qPCR directly targets DNA within durable oospores. This bypasses the high false-negative rates associated with physiological assays, delivering quantitative results from minimal soil volumes in a fraction of the time. Furthermore, while standard eDNA methods capture a mixture of intracellular and extracellular “relic” DNA, the oDNA protocol utilises size-selection filtration to enrich for oospores. This provides a high-fidelity enrichment of infectious propagule load rather than transient genetic signatures, enabling the detection of the shifts in pathogen load reported here.

In summary, our findings provide evidence-based guidance for optimal soil sampling and highlight the value of quantitative molecular monitoring. For routine surveillance, sampling the upper 10 cm optimises detection efficiency while minimising ecological impact on sensitive kauri roots. For high-priority monitoring, however, layered sampling (0–15 cm) could be used to provide insights into disease stage and/or total inoculum loads.

In addition, future work involving longitudinal monitoring of trees across disease stages— quantifying soil PA levels alongside visible health changes—could provide further valuable insights into infection thresholds, host responses, and PA persistence. For example, we have observed apparently healthy kauri (*e*.*g*., KM3) in this study and at other sites (unpublished data) with low PA levels (∼5 fg/g soil or less) and tracking these at-risk trees could reveal the PA concentration required for symptom development and identify potential resistance mechanisms. Similarly, monitoring trees with intermediate oospore loads (*e*.*g*., KM7, KM9) could provide insights into when oospore production peaks during the disease cycle, while tracking soil from around dead trees could determine PA persistence in soils.

Beyond *P. agathidicida*, this quantitative method is applicable to other *Phytophthora* species and research questions, offering a powerful tool for mapping horizontal spread, informing containment strategies (*e*.*g*., fencing), and evaluating the efficacy of treatments such as phosphite. By moving from qualitative presence/absence to high-resolution quantification, we can better understand and manage the small-scale pathogen dynamics that drive large-scale forest decline.

## ACKNOWLEDGEMENTS

We thank both the Kauri Ora Team of Te Roroa Iwi and BioSense for their assistance in collecting the soil samples used in this study. We also gratefully acknowledge Tiakina Kauri (part of Biosecurity New Zealand, a business unit of the Ministry for Primary Industries (MPI) for funding to support this research. Jade Palmer also acknowledges scholarship support from Tiakina Kauri, Victoria University of Wellington, the New Zealand Plant Protection Society, and Te Roroa Iwi. We thank Rena Misa for assistance in designing the initial version of Figure 4.

