## Supplemental Info for "Vertical distribution of *Phytophthora agathidicida* oospore DNA in kauri forest soils: Implications for optimised sampling and disease monitoring"

**Supplementary Figure 1**

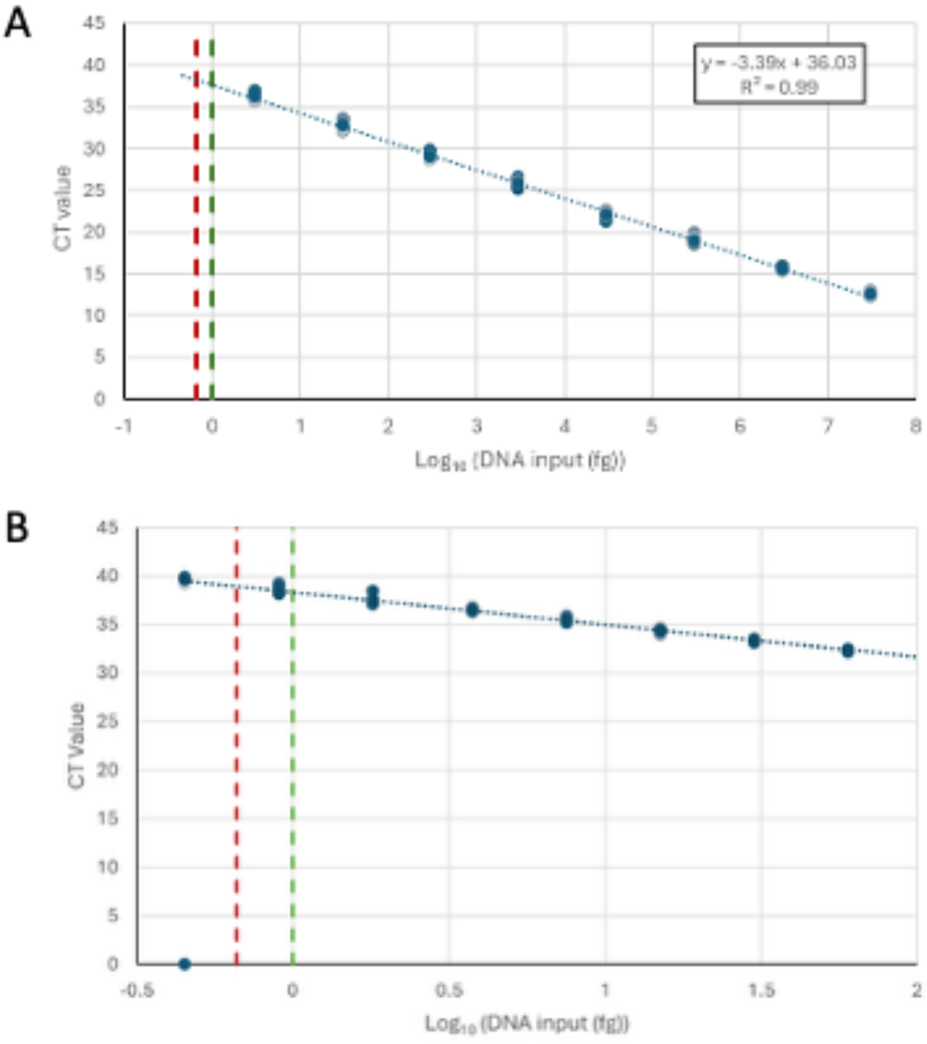


**Supplementary Figure 1. Performance and sensitivity of the Phytophthora agathidicida (PA) qPCR assay. (A) Standard curve and amplification efficiency.** qPCR standard curve generated using PA genomic DNA. Ten standards (ranging from 0.03 fg to 30,000,000 fg (30 ng) were measured using nine replicates per concentration. The lowest standard (0.03 fg) was not detected, and the second-lowest (0.3 fg) was detected in only two replicates and therefore excluded from curve fitting. Using the PA-LTR primers and the PA-LTR-probe, the assay exhibited an amplification efficiency of 97%. The upper limit of quantification was determined to be 30 ng, and the most linear range was 3,000,000 fg (3 ng) down to 3 fg. **(B) Limit of Detection (LoD) and Limit of Quantitation (LoQ).** To precisely define assay sensitivity, nine replicates of seven gDNA standards in twofold dilutions from 0. 16 fg to 60 fg were tested. LoD and LoQ values were calculated using algorithms optimised for eDNA assays (1). The coefficient of variation (CV) threshold for the LoQ was set to 0.35, and the 'Best' model settings were applied, as recommended. The LoD (red dashed line) was determined to be 0.7 fg, and the LoQ (green dashed line) was determined to be 1 fg. These thresholds delineate the technical limits for detecting and quantifying PA in environmental DNA extracts, distinguishing true low-level detections from stochastic amplification near the assay's limit.
